# A temporal sequence of heterochronic gene activities promotes stage-specific developmental events in *C. elegans*

**DOI:** 10.1101/2024.02.26.582127

**Authors:** Maria Ivanova, Eric G. Moss

## Abstract

The heterochronic genes of the nematode *Caenorhabditis elegans* control the succession of postembryonic developmental events. The four core heterochronic genes *lin-14, lin-28, hbl-1,* and *lin-41* act in a sequence to specify cell fates specific to each of the four larval stages. It was previously shown that *lin-14* has two activities separated in time that promote L1 and L2 developmental events, respectively. Using the auxin-inducible degron system, we find that *lin-28* and *hbl-1* each have two activities that control L2 and L3 events which are also separated in time. Relative to events they control, both *lin-28* and *hbl-1* appear to act just prior to or concurrently with events of the L2. Relative to each other, *lin-28* and *hbl-1* appear to act simultaneously. By contrast, the *lin-14* activity controlling L2 events precedes those of *lin-28* and *hbl-1* controlling the same events, suggesting *lin-14*’s regulation of *lin-28* is responsible for the delay. Likewise, the activities of *lin-28* and *hbl-1* controlling L3 fates act well in advance of those fates, suggesting a similar regulatory gap. *lin-41* acts early in the L3 to affect fates of the L4, although it was not possible to determine whether it too has two temporally separated activities. We also uncovered a feedback phenomenon that prevents the reactivation of heterochronic gene activity late in development after it has been down-regulated. This study places the heterochronic gene activities into a timeline of postembryonic development relative to one another and to the developmental events whose timing they control.

## Introduction

Postembryonic development of the nematode *Caenorhabditis elegans* occurs in four larval stages (L1-L4) where stage-specific developmental events—including cell divisions, differentiation, and morphogenesis—complete the formation of tissues and organs of the reproductive adult. The timing and sequence of these stage-specific events is under the control of the heterochronic genes, which act in a hierarchy that controls both the stage-specific events and the activities of later-acting heterochronic regulators (Rougvie and Moss, 2013). Four heterochronic genes, *lin-14, lin-28, hbl-1*, and *lin-41*, are the core factors specifying stage-specific events. Although their phenotypic effects and genetic relationships have been well-characterized, it is not known precisely when they act relative to each other and to the events they control. Outstanding questions are: Do genes that control the same cell fate events act at the same time, and do the genes act during the execution of the event they control, or prior to that event?

Ambros and Horvitz used temperature-sensitive alleles of *lin-14* to determine its times of action (Ambros and Horvitz, 1987). They shifted animals from the permissive to the restrictive temperature, and *vice versa*, at specific times during postembryonic development. These experiments showed that *lin-14* has two activities that occur at different times: an early activity, named *lin-14a*, that occurs early in the L1 stage that promotes L1-specific developmental events, and a second activity, *lin-14b*, that occurs hours later, in the middle of the L1 stage that promotes L2-specific events. The animal’s requirement for *lin-14a* occurs very close to the time the L1 events occur, whereas *lin-14b* occurs a few hours in advance of the L2 events that it controls (Ambros and Horvitz, 1987).

Similar studies have not been carried out for the other core heterochronic genes, *lin-28*, *hbl-1*, and *lin-41*, because temperature-sensitive alleles of these have not been identified. It is known that *lin-28* controls both L2- and L3-specific events, suggesting that it too has two separate activities (Ambros and Horvitz, 1984; Vadla *et al*., 2012). Like *lin-28* mutants, hypomorphic alleles of *hbl-1* also cause skipping of L2 events and precocious differentiation, but it has not yet been suggested that it has two activities (Abrahante *et al*., 2003, Abbott *et al*., 2005). *lin-41* controls events after *lin-28* and *hbl-1*, but it is unclear what stage *lin-41* acts on, whether L3 or L4 (Slack *et al*., 2000; Vadla *et al*., 2012).

To induce rapid degradation of these proteins without temperature-sensitive alleles, we used the auxin-inducible degron (AID) system adapted from *Arabidopsis thaliana* (Zhang *et al*. 2015, Hills-Muckey *et al*., 2022). We constructed alleles of each of these genes with the degron coding region fused to either the 5’- or 3’- end of a gene’s open reading frame using CRISPR/Cas9. All of these genes were wildtype in phenotype in the absence of auxin analog when we used a modified TIR (*TIR1(F79G)*; Hills-Muckey *et al*., 2022). This approach allowed us to determine the times of action of these heterochronic genes and their temporal relationships to each other and the events they control.

## Results

### The times of actions for *lin-14* determined by *AID* match those determined by temperature-sensitive alleles

To test the usefulness of the auxin-inducible degron system for determining the times of action of the heterochronic genes, we analyzed the phenotypes of *lin-14::AID* animals transferred to and from the auxin analog 5-Ph-IAA during larval development. Shifting animals onto 5-Ph-IAA should be equivalent to shifting temperature-sensitive (*ts*) alleles to the non-permissive temperature.

We first studied *lin-14*’s control of L2 events (the *lin-14b* activity) by assessing seam cell number at adulthood of animals shifted at different times. Normally, the total number of seam cells increases during the L2 due to a stage-specific symmetric division in many of these cells (Sulston and Horvitz, 1977). If the L2-specific seam cell lineage patterns are skipped, and L3 patterns occur precocious in their place due to the lack of *lin-14b* activity, adult seam cell numbers will be reduced (Ambros and Horvitz, 1987).

We observed that the number of seam cells at adulthood was the lowest when *lin-14::AID* animals were shifted to 5-Ph-IAA at 4-8 hours after hatching (Fig. 1A). Larvae transferred to 5-Ph-IAA at 10 and 12 hours had an intermediate number of seam cells, and most larvae executed normal seam cell division patterns when they were transferred to 5-Ph-IAA at 14 hours of postembryonic development or later (Fig. 1A).

**Fig. 1.**
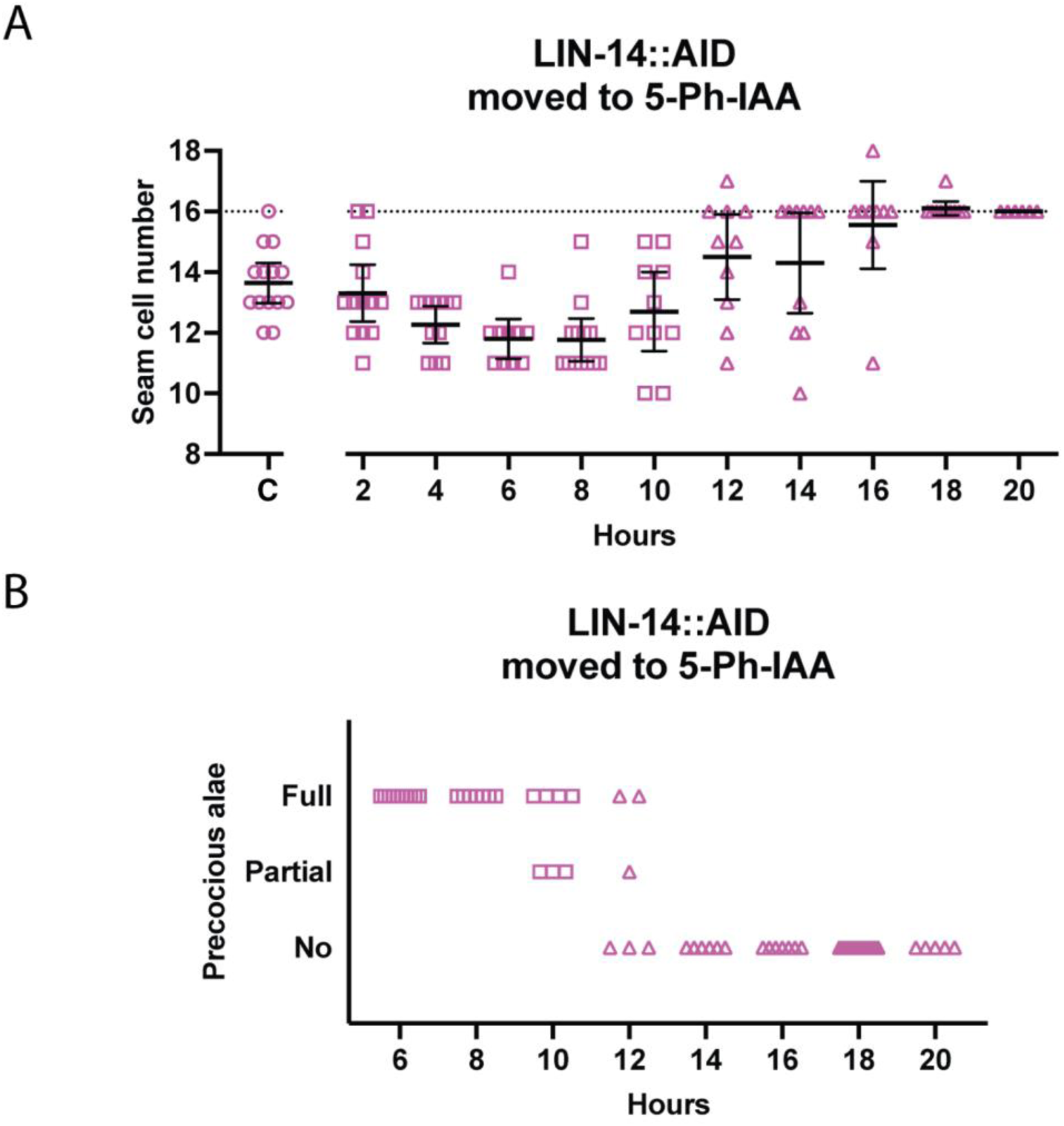
lin-14 acts during the mid-L1 stage to promote L2 seam cell fates. (A) Numbers of seam cells in lin-14::AID adults moved to 5-Ph-IAA at the indicated times after synchronization. Horizontal dotted line indicates the wildtype number of seam cells, 16. ‘C’ indicates animals grown continuously on 5-Ph-IAA. Bars indicate averages and 95% CI. (B) Precocious adult alae in lin-14::AID animals moved to 5-Ph-IAA at the indicated times after synchronization. Alae data was recorded only for L4 larvae. Different datapoint shapes indicate different groups of synchronized animals; that were necessary to cover all the time points. Several animals at 14 hours had fewer than 16 seam cells, those animals also were at the late L3 stage and the alae data is not known for them and not shown in the second plot. The L1 lethargus began between 16 and 18 hours.

Ambros and Horvitz considered animals with 30% or fewer seam cells expressing L3 patterns equivalent to those grown at the permissive temperature (Ambros and Horvitz, 1987). In our experiments, 70% of worms transferred to 5-Ph-IAA at 12 hours had 14 or more seam cells, meaning that they had fewer than 30% of seam cells expressing L3 patterns. Thus, the 12-hour time point is the end of *lin-14*’s auxin-sensitive period, which closely corresponds to the period defined using temperature-sensitive alleles.

Next, we examined the formation of adult alae. Adult alae are cuticle structures that form when seam cells differentiate at adulthood. Defective *lin-14b*, like other heterochronic mutants that result in precocious development, cause adult alae to form at least one stage early (Ambros and Horvitz, 1984; Ambros and Horvitz, 1987; Ambros, 1989). It is not believed that *lin-14* directly controls alae formation, rather that the time of alae formation is a consequence of *lin-14*’s activity earlier, specifically on L2 events. *lin-14::AID* larvae transferred to 5-Ph-IAA at 6 hours had full precocious alae, but when transferred between 10 and 12 hours in development, they displayed partial or no precocious alae, and alae developed at the normal time when they were transferred to 5-Ph-IAA at 14 hours in development or later (Fig. 1B).

For most *lin-14::AID* animals grown with or without 5-Ph-IAA, L1 seam cell divisions occurred at 4-7 hours, and animals entered lethargus at the end of L1 between 16-18 hours of larval development. Thus, the end of the time period requiring *lin-14b* activity as determined by the AID system (its auxin-sensitive period) was the mid-to late-L1 stage, which matches the temperature-sensitive period of *lin-14b* defined using temperature-sensitive alleles (Ambros and Horvitz, 1984).

*lin-14a* activity specifies developmental events of the L1, and lack of *lin-14a* causes L2 events to be executed in the L1 (Ambros and Horvitz, 1987). Because *lin-14(0)* animals lack both *lin-14a* and *lin-14b* activities, they execute the L2-specific symmetric division of seam cells only once, precociously in the L1, and thereby end up with a normal adult seam cell count, despite other problems (Ambros and Horvitz, 1987). It is important to note that we found that *lin-14::AID* animals grown continuously on 5-Ph-IAA did not have a normal count but rather had a reduced number of seam cells, indicating that *lin-14a* activity was not completely reduced to null levels (Fig. 1A). The degron, therefore, did not appear to completely inactivate *lin-14,* but rather resulted in a partial loss-of-function instead of a null phenotype. Nevertheless, we could still use this condition to characterize *lin-14a* timing.

Shifting *lin-14::AID* animals away from 5-Ph-IAA is the equivalent of shifting *ts*-alleles to the permissive temperature. Most *lin-14::AID* larvae removed from 5-Ph-IAA in the first hour in development had a higher than normal number of seam cells at adulthood but no precocious alae (Fig. 2). The same phenotype was observed by Ambros and Horvitz (1984) as a result of the L2 fates being executed twice, in both the L1 and L2, and interpreted as a loss of *lin-14a* activity with normal *lin-14b* activity (*lin-14a^−^b^+^*). Shifting animals at 2 hours resulted in a nearly wild-type level of seam cells and some precocious alae. These data indicate that the auxin-sensitive period for the *lin-14a* activity begins at 1 hour or before and that the auxin-sensitive period for the *lin-14b* begins around 2 hours.

**Fig. 2.**
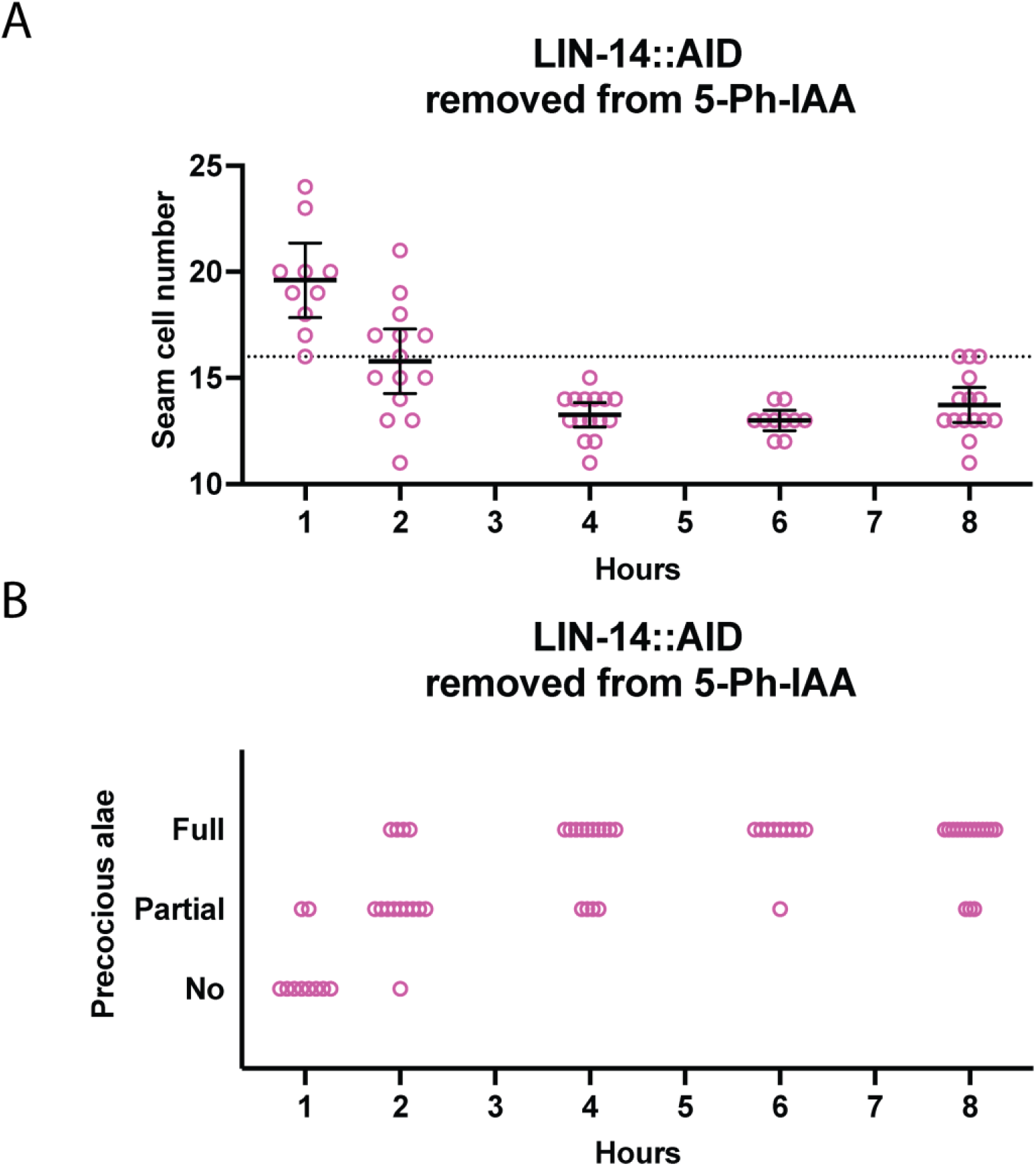
lin-14 activity controlling L1 seam cell fates occurs prior to 2 hours of postembryonic development. (A) Numbers of seam cells in lin-14::AID animals removed from 5-Ph-IAA at the indicated times after synchronization. Horizontal dotted line indicates the exact wild-type number of seam cells. Bars indicate averages and 95% CIs. (B) Precocious adult alae in lin-14::AID animals removed from 5-Ph-IAA at the indicated times after synchronization. Alae data was recorded only for L4 larvae.

Although we were not able to observe a time frame for the restoration of *lin-14a* activity, *lin-14b* activity could be fully restored when worms were removed from 5-Ph-IAA plates at 1 hour after start, and at 2 hours in some worms, whereas Ambros and Horvitz observed restoration following shifts up until 6-7 hours in development (Ambros and Horvitz, 1987). Thus, the recovery from 5-Ph-IAA is not as rapid as the forward experiments for *lin-14*. But overall, these results showed that the AID system could be used to determine the times of actions for heterochronic genes and that times obtained in forward experiments are precise.

Intestinal nuclei divisions occur during the L1 stage, and these are also under the control of *lin-14*. *C. elegans* L1 animals hatch with 20 intestinal nuclei, but some of these undergo mitosis during the L1 molt to yield 30-34 intestinal nuclei by adulthood. Our own counts of the *lin-14::AID* strain showed that early L1 larvae had 20 ± 0.5 (n = 14) intestinal nuclei, and animals after L2 seam cell divisions had 30 ± 2.3 (n = 12) intestinal nuclei. Consistent with prior observations of *lin-14(0)* mutants, we observed that *lin-14::AID* animals grown continuously on 5-Ph-IAA had only 21.8 ± 2.4 (n = 15) intestinal nuclei at adulthood in contrast to normal 30-34 (Sulston and Horvitz, 1977; Ambros and Horvitz, 1984).

We determined the auxin-sensitive period for *lin-14*’s control of intestinal nuclear divisions. Intestinal nuclei counts were reduced in *lin-14::AID* animals transferred to 5-Ph-IAA before 16 hours of postembryonic development. L1 lethargus was observed at 16-18 hours, and the animals in lethargus (without pharyngeal pumping) that were transferred to 5-Ph-IAA at 18 hours developed a normal number of intestinal nuclei (Fig. 3). The number of intestinal nuclei increased in animals transferred to 5-Ph-IAA at 13-14 hours in development as compared to those always grown on 5-Ph-IAA, yet was still less than the wild-type number, indicating that *lin-14* activity promoting intestinal nuclei divisions was not completed at that time. Thus, this activity of *lin-14* in the intestine has a slightly different time frame compared to its activities promoting seam cell fates. As a result, although the control of intestinal nuclei divisions and the *lin-14a* activity both occur in the L1 stage, these activities have different time frames and might be independent.

**Fig. 3.**
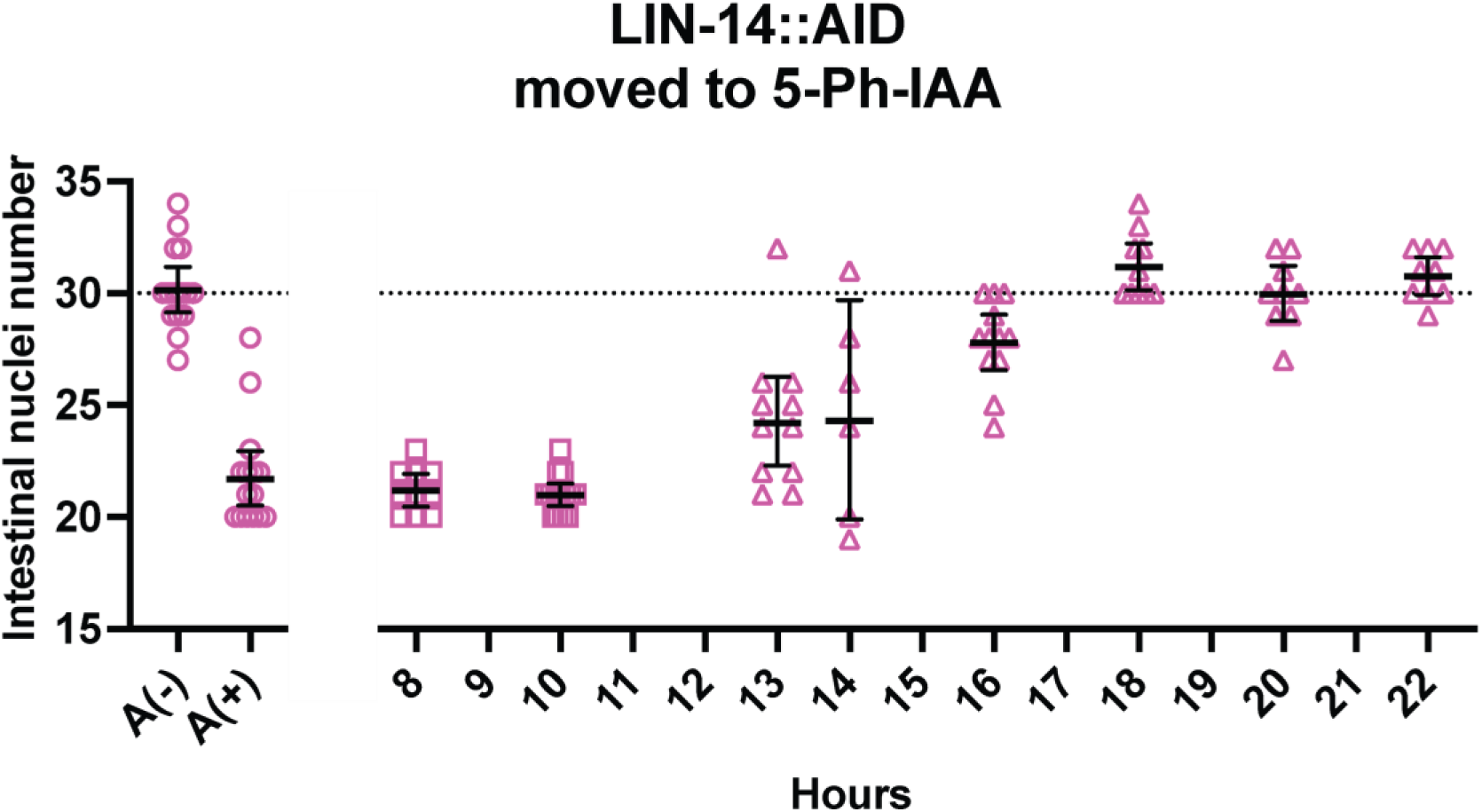
lin-14 activity promotes intestinal nuclei divisions prior to 16 hours of postembryonic development. Intestinal nuclei numbers in lin-14::AID animals moved to 5-Ph-IAA at the indicated times after synchronization. The horizontal dotted line indicates the average number of intestinal nuclei in wild-type animals. A(-) indicates animals grown continuously without 5-Ph-IAA. A(+) indicates animals grown continuously on 5-Ph-IAA. Bars indicate averages and 95% CIs. Different datapoint shapes indicate different groups of synchronized animals needed to cover all the time points.

### *lin-28 acts later than lin-14* to promote L2 developmental events

Having established for *lin-14* that the auxin-sensitive periods match the established temperature-sensitive periods, we employed this system to study *lin-28* and *hbl-1*.

*lin-28* mutants are like *lin-14b* mutants in that they skip L2-specific events, including the L2-specific symmetric seam cell division that increases the number of seam cells in late stages. (Ambros and Horvitz, 1984; Ambros and Horvitz, 1987; Seggerson et al., 2002). When *lin-28:AID* animals are grown continuously on 5-Ph-IAA, seam cell counts are reduced compared to wildtype (Fig. 4A). This reduced count is true for animals shifted onto 5-Ph-IAA from 8 to 14 hours. However, the counts increased near the wildtype level in animals transferred to 5-Ph-IAA from 16 to 20 hours of development (Fig. 4A). In general, wildtype seam cell counts were observed when animals were shifted at 22 hours or later; however, some *lin-28::AID* animals transferred to 5-Ph-IAA at 22 hours and later had 17 seam cells, the reason is unclear (Fig. 4A, Fig. 5A). Thus, *lin-28’s* time of action for specifying L2 cell fates as determined by AID was at the end of the L1 stage (Fig. 4A, Fig. 5A). Note that our experiments directly compared the auxin-sensitive periods of *lin-14* and *lin-28* by shifting both strains in parallel at the same time points (Table S1). We found that this period for *lin-28* was significantly later than what we determined for *lin-14b* activity, the mid-L1 stage, so that these genes do appear to be acting separately, not simultaneously, to specify L2 fates.

**Fig. 4.**
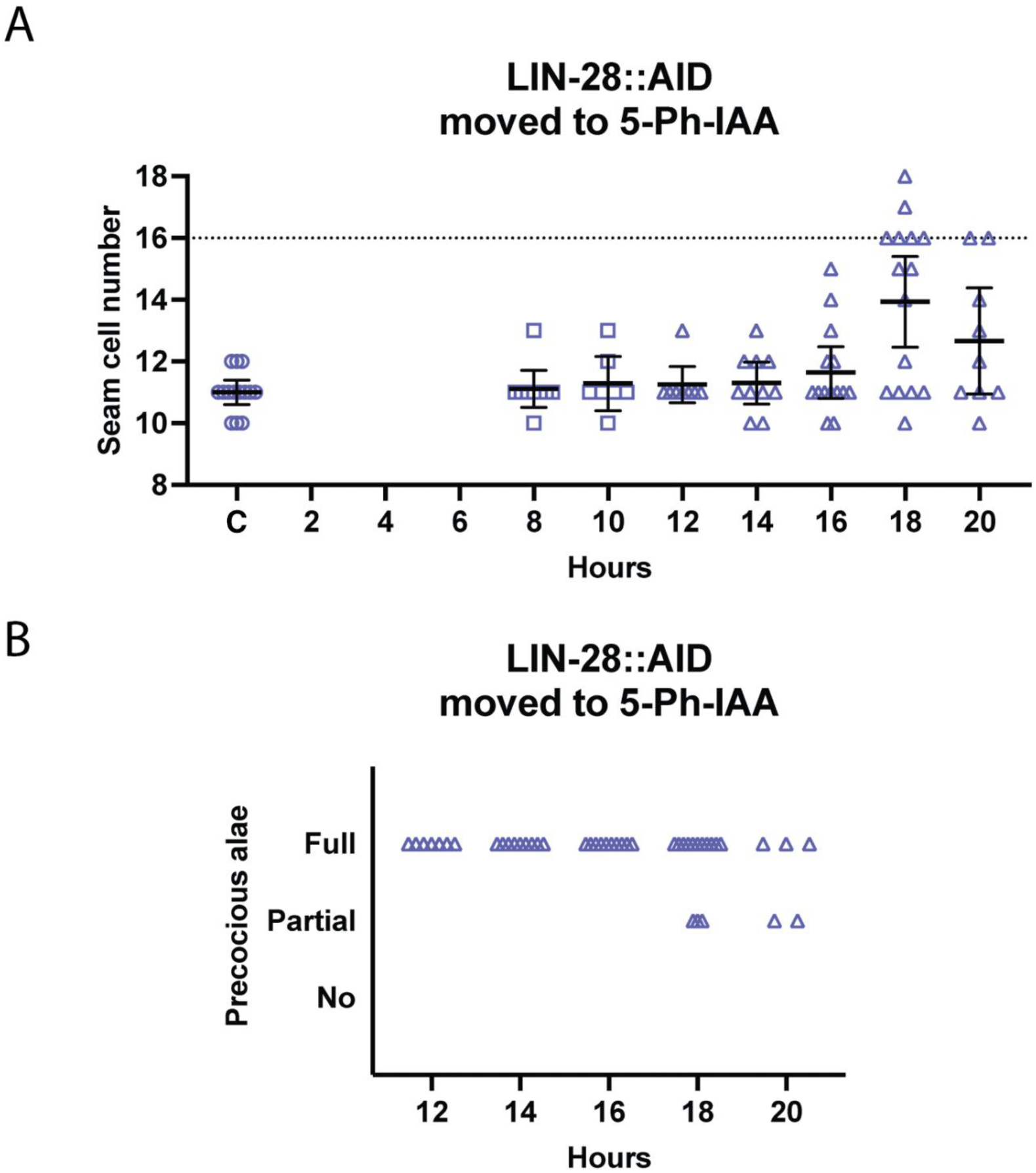
The activity of lin-28 that promotes L2 seam cell fates occurs later than the similar activity of lin-14. (A) Numbers of seam cells in lin-28::AID animals moved to 5-Ph-IAA at the indicated times after synchronization. The horizontal dotted line indicates the wild-type number of seam cells. ‘C’ indicates animals grown continuously on 5-Ph-IAA. Bars indicate averages and 95% CIs. (B) Adult alae in lin-28::AID animals moved to 5-Ph-IAA at the indicated times after synchronization in the same experiments shown in panel A. Alae data was recorded only for L4 larvae. Different dot shapes indicate different groups of synchronized animals.

**Fig. 5.**
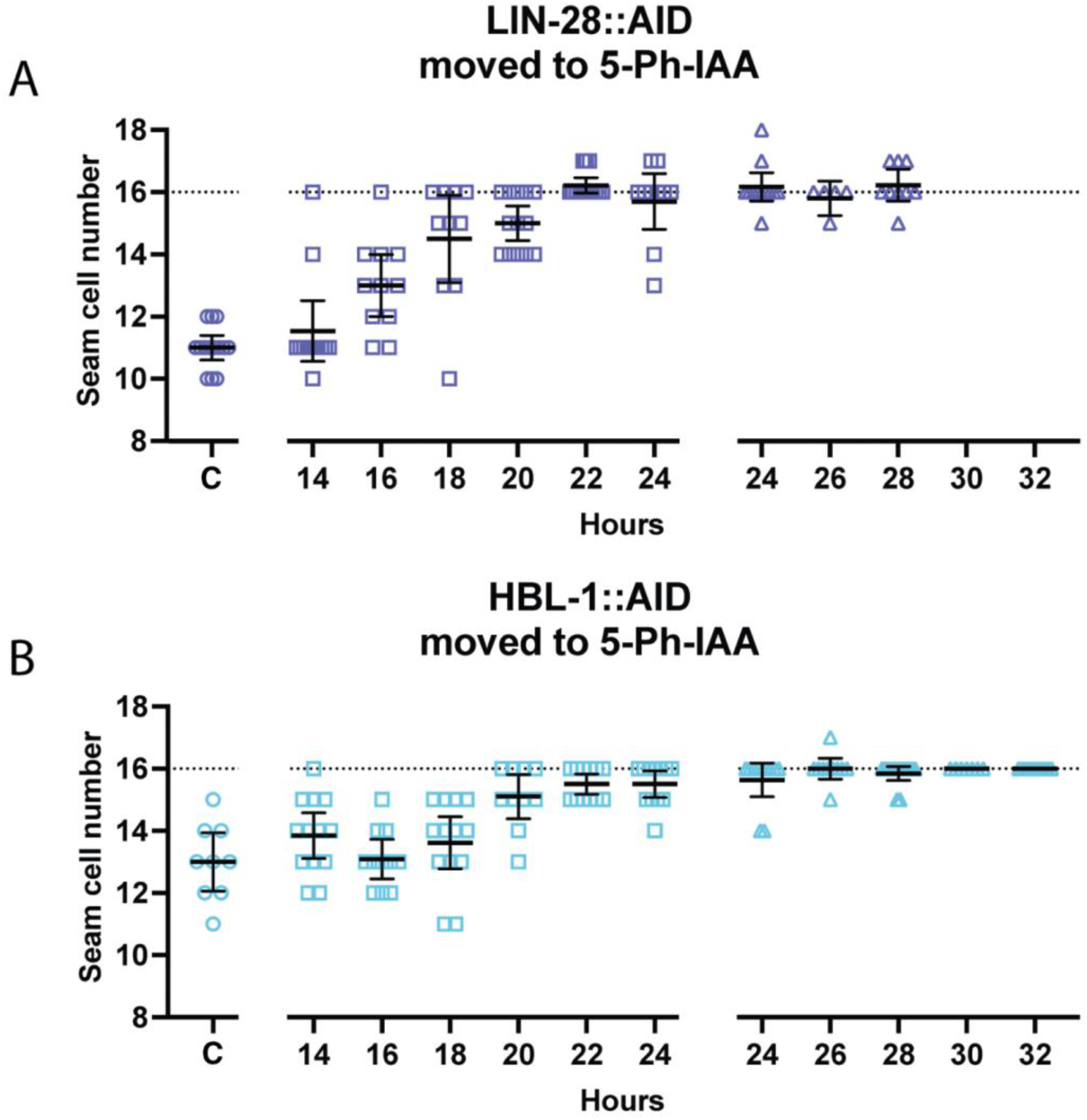
The activities of lin-28 and hbl-1 that promote L2 seam cell fates coincide in time. (A) Numbers of seam cells in lin-28::AID animals moved to 5-Ph-IAA at the indicated times after synchronization. (B) Numbers of seam cells in hbl-1::AID animals moved to 5-Ph-IAA at the indicated times after synchronization. Horizontal dotted lines indicate the wild-type number of seam cells. ‘C’ indicates animals grown continuously on 5-Ph-IAA. Bars indicate averages and 95% CIs. Different dot shapes indicate different groups of synchronized animals.

### lin-28 and hbl-1 act simultaneously to control L2 cell fates, just prior to L2 seam cell divisions

Loss-of-function alleles of *hbl-1* cause a phenotype similar to that of *lin-28* mutants (Abrahante *et al*., 2003). We directly compared the auxin-sensitive periods of these two genes (Table S1). The time frame for the end of *hbl-1* activity promoting L2 seam cell fates was almost identical to that of *lin-28* (Fig. 5). The majority of *hbl-1::AID* animals developed a normal number of seam cells when they were moved to 5-Ph-IAA at 24 hours in development or later (Fig. 5B).

The times of action of *lin-28* and *hbl-1* determined by AID appeared to coincide with the L1 lethargus that precedes the L1 molt. The symmetric seam cell divisions normally occur soon after the L1 molt (Sulston and Horvitz, 1977). Considering the time needed for the process of degradation to start working, we conclude that the activities of *lin-28* and *hbl-1* are required right before the L2 seam cell divisions.

### lin-28 and hbl-1 act during the L2 to promote later events

*lin-28*’s effect on the time of adult alae formation is understood to be a consequence of its control of L3 fates, which is a separate activity from its control of L2 fates (Vadla et al., 2012). For both *lin-28* and *hbl-1*, the auxin-sensitive period for precocious adult alae development was slightly later than that of the L2-specific seam cell increase (Figs. 4-6). Some animals transferred to 5-Ph-IAA at 20, 22, and 24 hours had normal numbers of seam cells and complete or gapped precocious alae (Fig. 6), which indicates that they executed the L2 seam cell fates normally and then skipped a later stage. Thus, like *lin-14*, both *lin-28* and *hbl-1* appear to have two activities separated in time.

**Fig. 6.**
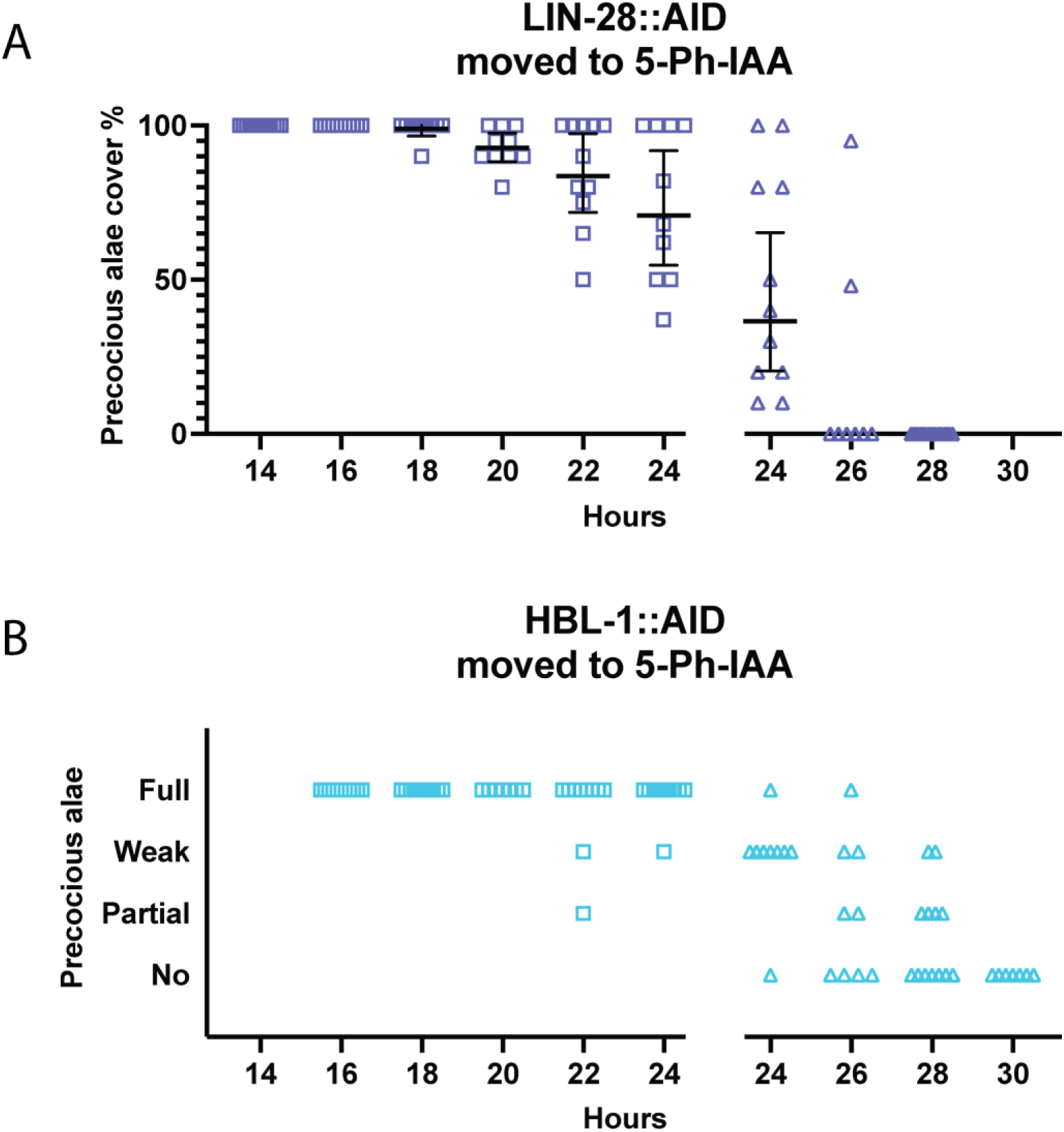
lin-28 and hbl-1 have activities that promote seam cell fates after the L2 stage. (A) Precocious alae in lin-28::AID animals moved to 5-Ph-IAA at the indicated times after synchronization. The percent was calculated as the fraction of seam cells that developed alae versus the total number of seam cells. Bars indicate averages and 95% CIs. (B) Precocious alae in hbl-1::AID animals moved to 5-Ph-IAA at the indicated times after synchronization. Alae were scored as “full” when they could be easily followed from head to tail, “weak” when alae did not have clear gaps but also could not be followed completely, and “partial” when gaps were observed. A gap in the graph separates different experiments. Different dot shapes indicate different groups of synchronized animals.

We noted a difference in the appearance of the precocious alae of *lin-28::AID* and *hbl-1::AID* strains (Fig. S1). In *lin-28::AID* animals, precocious alae were strong even when they had gaps, whereas the *hbl-1::AID* alae were difficult to see because they were less pronounced or thinner. In most cases, it was difficult to estimate an approximate percent expressivity of the precocious alae in these animals. This observation suggests some difference in how *lin-28* and *hbl-1* affect the differentiation of seam cells.

### *lin-28* is *subject to* a feedback loop

In experiments where we shifted *lin-28:AID* animals from 5-Ph-IAA to plates without the auxin analog, we found that *lin-28* activity did not recover. Specifically, L1 larvae that were placed on 5-Ph-IAA after synchronization and removed in 1 hour developed a null-like phenotype with reduced numbers of seam cells and precocious alae (11 ± 0.5 seam cells, 87% with full precocious alae, n = 8). This was surprising given the fact that experiments shifting these animals onto 5-Ph-IAA showed that the auxin-sensitive period for *lin-28* extended well past 1 hour of postembryonic development. This finding suggests a feedback loop: once *lin-28* activity is reduced, consequences of its downregulation make it impossible either for *lin-28* expression to be restored or for *lin-28* to have an effect on downstream events. This phenomenon prevented us from defining a precise start time of *lin-28*’s activity in the L1.

### *hbl-1* has two activities separated in time

In contrast to *lin-28*, *hbl-1*’s activity was restored when shifted away from the auxin analog 5-Ph-IAA. When synchronized *hbl-1::AID* animals were grown on 5-Ph-IAA from the start of postembryonic development and then transferred away at 4 hour intervals, both seam cell numbers and adult alae formation were mostly wild-type up until 20 hours, indicating recovery from the degradation over that timeframe (Table S1; Fig. 7). It was not until the *hbl-1::AID* animals were on 5-Ph-IAA for 24 hours or longer did precocious phenotypes occur (Fig. 7).

**Fig. 7.**
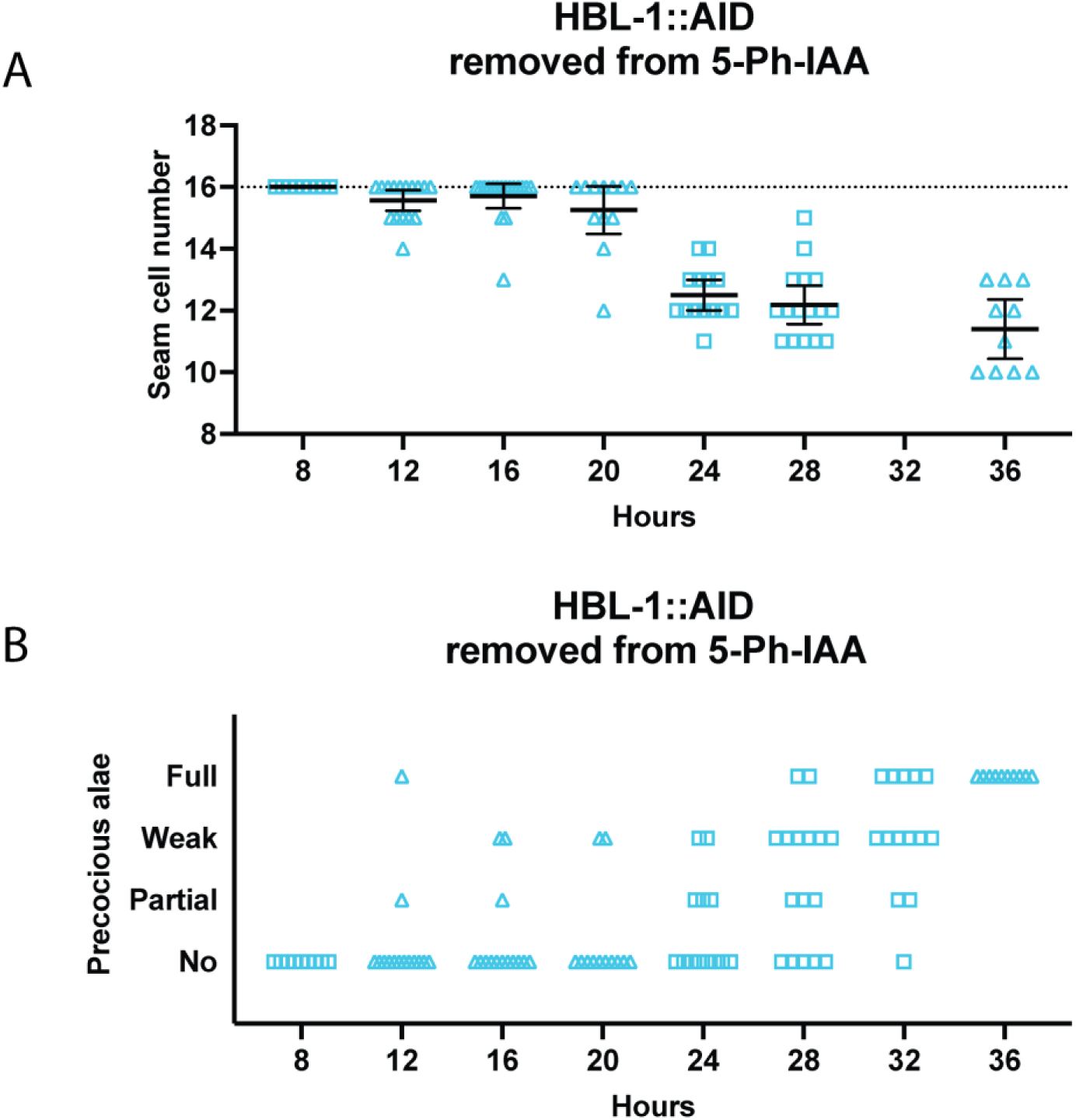
hbl-1 activity is restored after removal from 5-Ph-IAA. (A) Numbers of seam cells in hbl-1::AID animals removed from 5-Ph-IAA at the indicated times after synchronization. Horizontal dotted lines indicate the wild-type number of seam cells. Bars indicate averages and 95% CIs. (B) Precocious alae in hbl-1::AID animals removed from 5-Ph-IAA at the indicated times after synchronization. Different dot shapes indicate different groups of synchronized animals.

To define the gene’s auxin-sensitive period for determining L2 seam cell fates, *hbl-1::AID* animals were either transferred to 5-Ph-IAA or removed from it in 2-hour intervals, after which both seam cell counts and precocious alae formation were assessed (Table S1; Fig. 8). Experiments where animals were shifted to 5-Ph-IAA showed that the auxin sensitive period is over by about 22 hours of postembryonic development, as seam cell counts were mostly wild-type from that point on (Fig. 8A). In experiments where *hbl-1::AID* animals were moved off the auxin analog, seam cells remained normal in animals removed from 5-Ph-IAA before 18 hours of larval development and showed a strong loss-of-function phenotype in those removed at 24 hours or later (Fig. 8B). These results define *hbl-1*’s auxin-sensitive period to between 18-22 hours of post-embryonic development. We observed L2 seam cell divisions to occur between 21-24 hours in *hbl-1::AID* animals grown either without or continuously on 5-Ph-IAA. Therefore, *hbl-1*’s time of action appears to occur just before or during the L2 seam cell divisions.

**Fig. 8.**
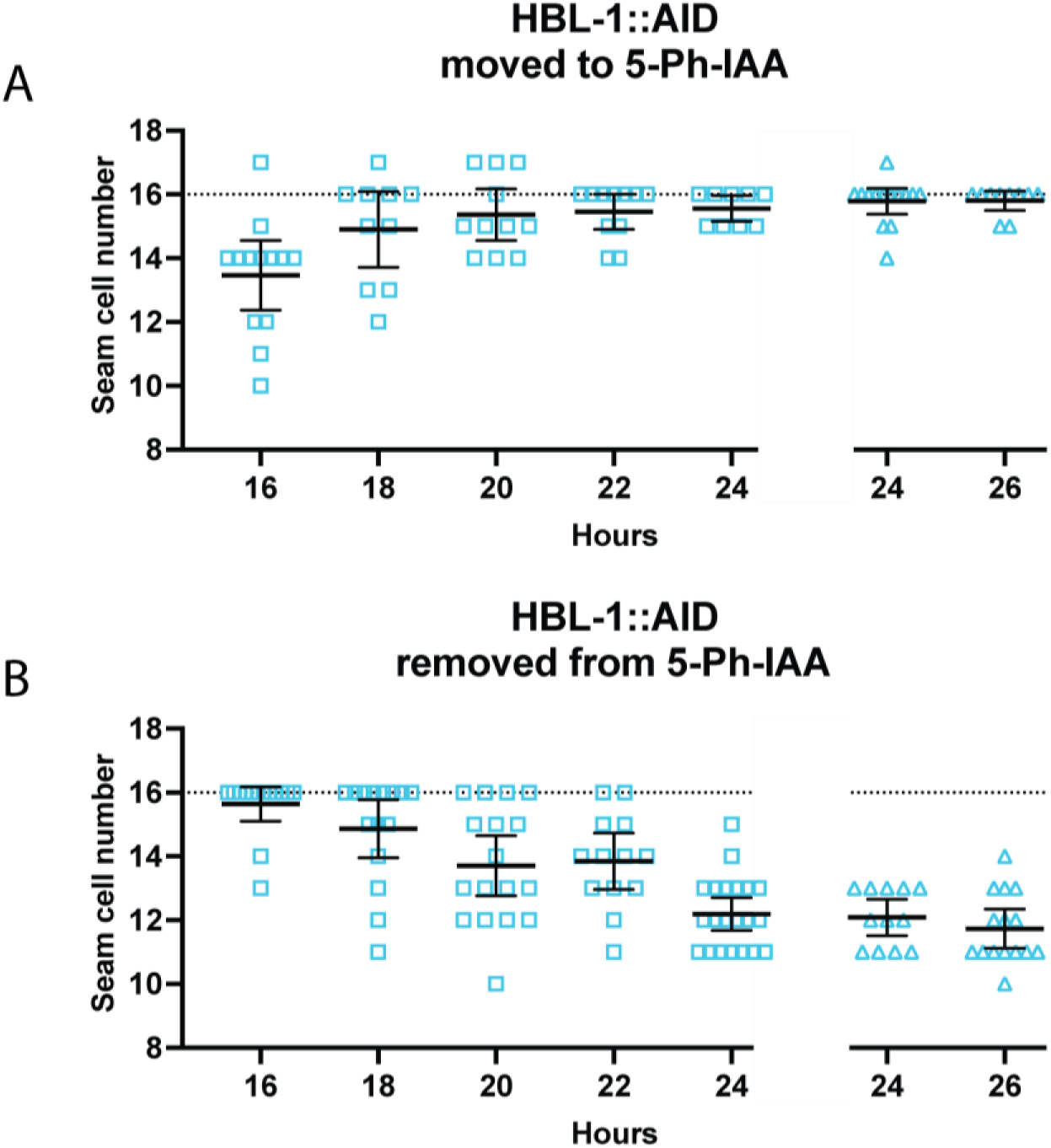
hbl-1 acts to promote L2 seam cell fates over a narrow timeframe. (A) Numbers of seam cells in hbl-1::AID animals transferred to 5-Ph-IAA at the indicated times after synchronization. (B) Numbers of seam cells in hbl-1::AID animals removed from 5-Ph-IAA at the indicated times after synchronization. Horizontal dotted lines indicate the exact wild-type number of seam cells. Bars indicate averages and 95% CIs. Gaps in the graphs separate different experiments. Different dot shapes indicate different groups of synchronized animals.

Because *lin-28*’s effect on the L2-specific division pattern and seam cell differentiation at adulthood reflect two different activities of the same gene, we wished to assess whether the same is true for *hbl-1* (Vadla, et al. 2012). We observed that full precocious alae appeared in *hbl-1::AID* animals that had been transferred to 5-Ph-IAA before 24 hours in development. The development of precocious alae became less frequent in later transfers, until they disappeared altogether in animals transferred at 30 hours in development (Fig. 9A). Similarly, in animals removed from 5-Ph-IAA at 24-26 hours in development, precocious alae sometimes appeared (Fig. 9B). Since most animals did not develop precocious alae when they were moved to 5-Ph-IAA at 28 hours or later, and most animals formed precocious alae if they were removed from 5-Ph-IAA at the same time, this must be shortly after the time that *hbl-1* acts to control later events. Thus, *hbl-1*’s activity promoting seam cell fates after the L2 (24-26 hours) is separated in time from its earlier activity controlling seam cell divisions (18-22 hours). This second activity ceases in 4-6 hours after the L2 divisions are completed, which is approximately 4-6 hours before the L2 molt.

**Fig. 9.**
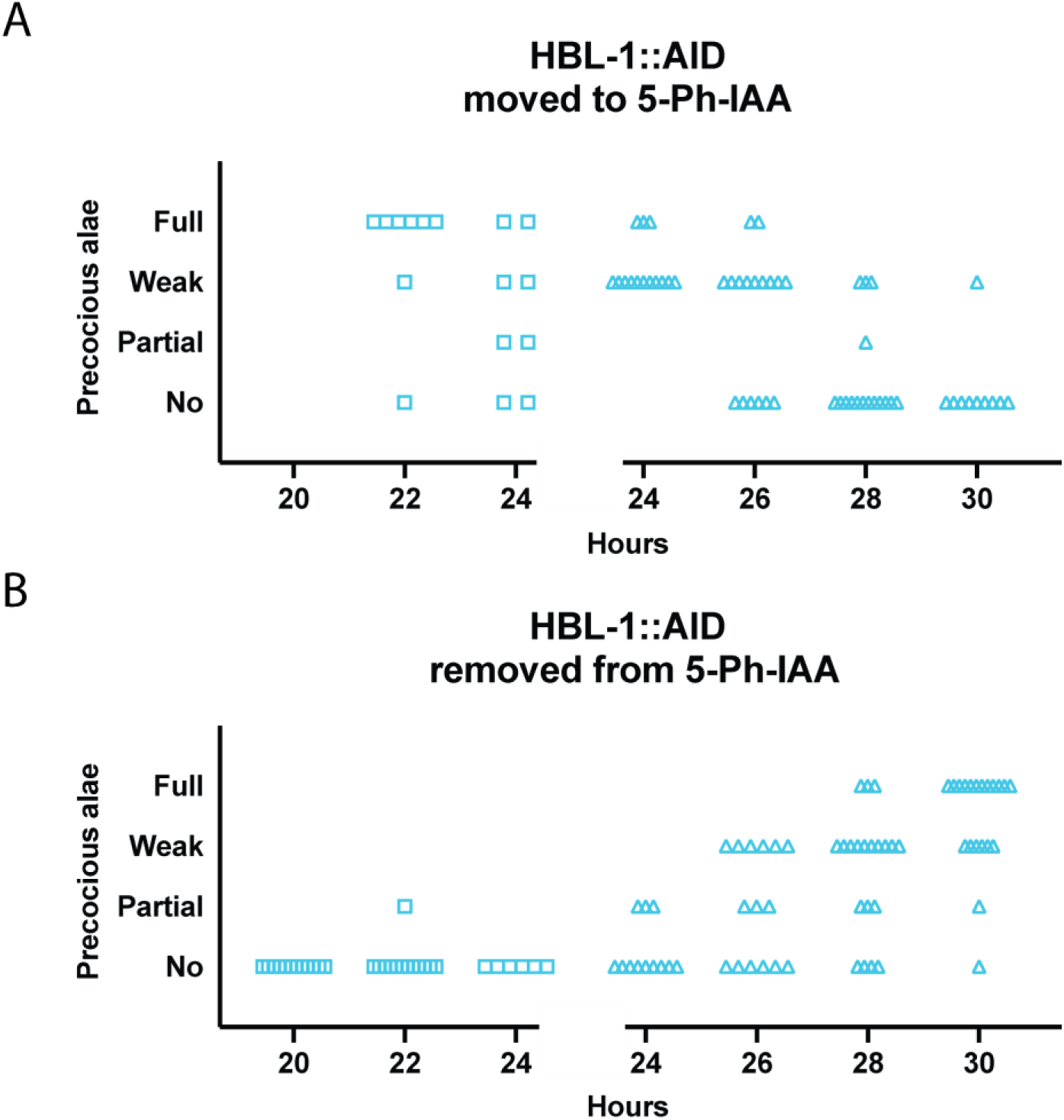
The time of the second hbl-1 activity is distinct from the first. (A) Precocious alae in hbl-1::AID animals moved to 5-Ph-IAA at the indicated times after synchronization. (B) Precocious alae in hbl-1::AID animals removed from 5-Ph-IAA at the indicated times after synchronization. Alae were scored as “full” if they could be easily followed from head to tail, “weak” if they did not have clear gaps but could not be easily followed, and “partial” when gaps were observed. Gaps in graphs separate different experiments. Different dot shapes indicate different groups of synchronized animals.

### hbl-1 deficiency causes skipping of L3 developmental events

Hermaphrodite seam cell lineages show the same division patterns in L3 and L4 stages, so it is not possible to distinguish skipping of L3 versus L4 cell fates in these lineages. However, certain male seam cell lineages, those that give rise to the male tail rays, do have pattern differences between these stages (Sulston and Horvitz, 1977).

Like the seam cells in the L2, 5 male seam cells undergo additional symmetric division in the L3 that will ultimately produce 9 ray precursor cells that will form the rays of the male tail (Sulston and Horvitz, 1977). We predicted that if the L3-specific fates were skipped there would be a reduced number of ray precursor cells; if the L4 fates were skipped, the number of ray precursor cells would not be reduced.

First, we grew *hbl-1::AID* animals continuously on 5-Ph-IAA and examined mail tails between 32-50 hours when ray cell divisions occur. We observed that the number of ray precursor cells was significantly reduced and ray differentiation occurred precociously: at 32 hours, ray cells already completed L4 specific divisions, while in animals grown on plates without auxin they were still undergoing L3 specific divisions (n=15, Fig. 10). Note that in addition to occurring in the L2, symmetric seam cell divisions also occur in the L3 in ray-producing seam cells V6 and V7 (Sulston and Horvitz, 1977). So, the reduction of ray precursor cells in *hbl-1::AID* animals grown continuously on the auxin analog could be due to loss of these proliferative divisions in both L2 and L3.

**Fig. 10.**
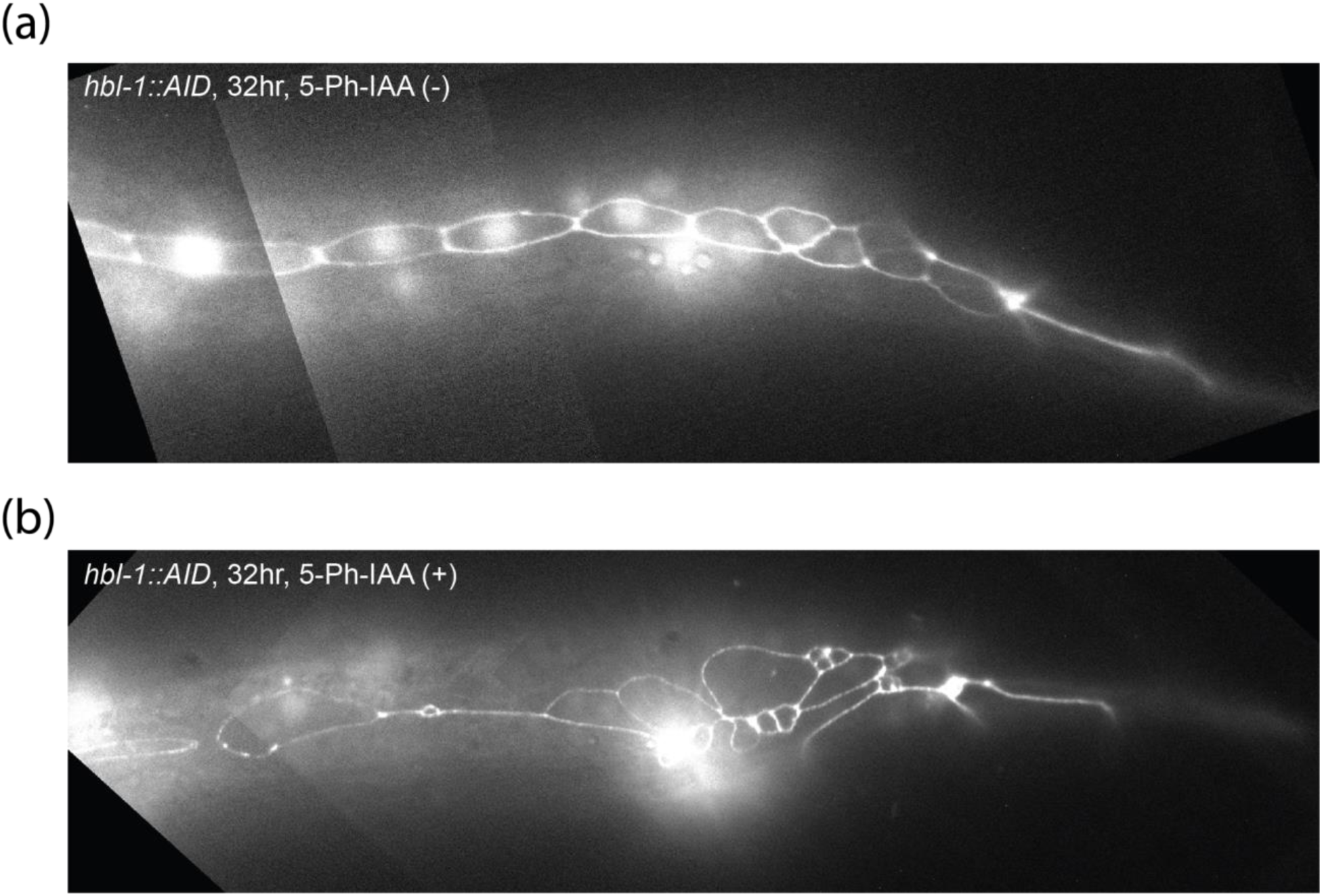
Reduction of hbl-1 activity causes a skipping of L3 stage and premature cell fates in male tail. (a) Fluorescence micrograph of a male tail at 32 hr in a hbl-1::AID animal grown without 5-Ph-IAA. The number of ray precursor cells appears reduced; however, those cells proceeded to undergo further divisions characteristic of these cells. (b) Fluorescent micrograph of male tail at 32 hr in a hbl-1::AID animal grown continuously on 5-Ph-IAA. Seam cell nuclei and cell junctions visualized with SCM::GFP and ajm-1::GFP, respectively. Animals are oriented anterior to the left, dorsal side up.

Next, *hbl-1::AID* animals were moved to auxin between 18 and 22 hours of postembryonic development (Fig. 11). We examined both the non-ray producing seam cells, V1-V4, and the ray-producing seam cells V5, V6, and T. For each time point, 7 animals were observed, all animals had patterns similar to those represented on micrographs in Fig. 11. Animals moved to auxin at either 18 or 20 hours showed precocious differentiation of precursor cells and a reduced number of rays, but had a reduced overall number of seam cells (12-13), meaning the reduction in rays may have been due, at least in part, to skipping the symmetric division of the L2. In contrast, animals moved to auxin at the slightly later time point of 22 hours of development showed reduced number of rays but overall the wildtype number of seam cells (16) (Fig. 11), meaning the L2-specific symmetric division was not skipped. We interpret these observations to mean that the reduction in the second of *hbl-1*’s two activities can cause the skipping of L3 specific divisions to lead to precocious L4 and adult cell fates.

**Fig. 11.**
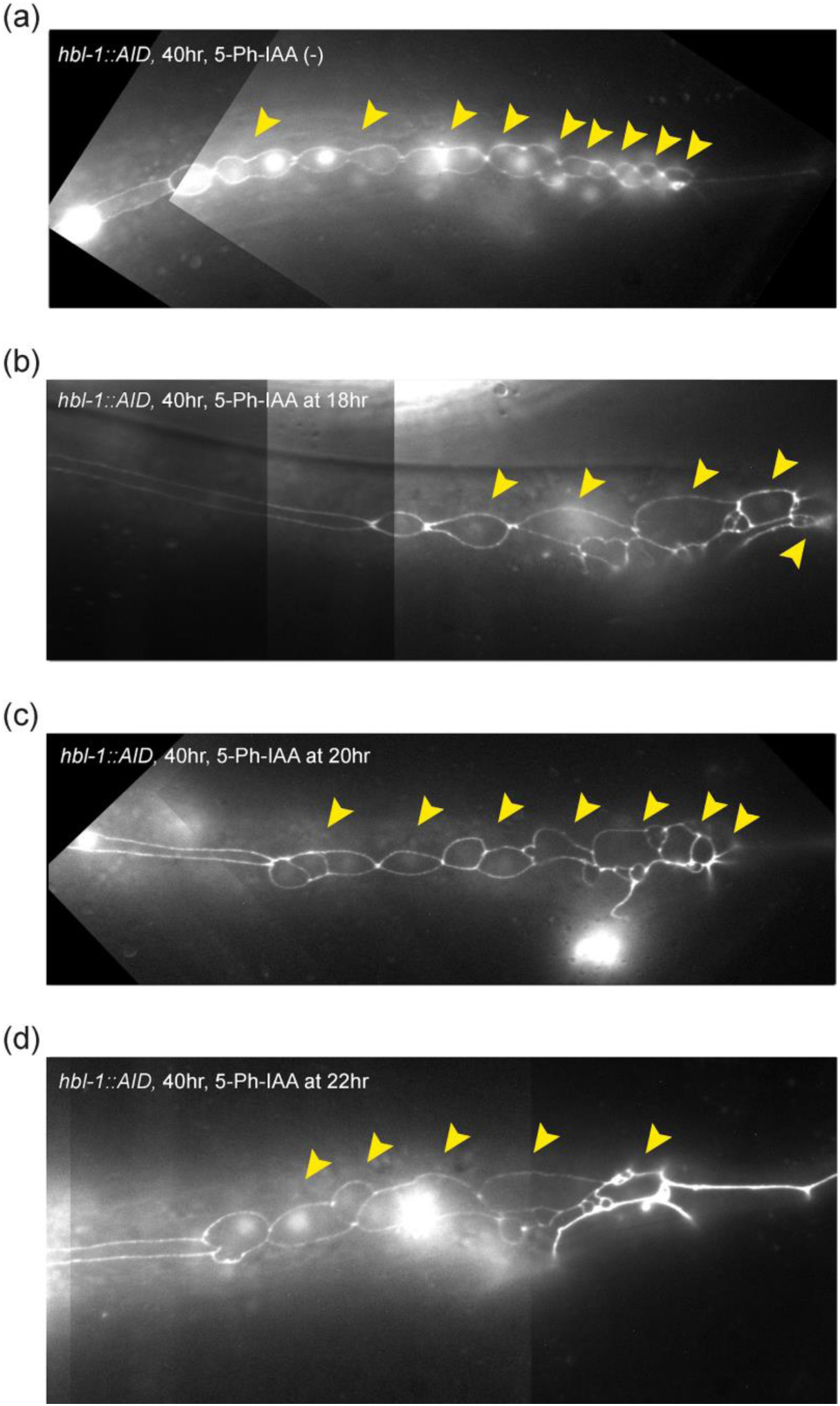
HBL-1 degradation at certain times in development causes a skipping of L3 stage and premature cell fates in male tail. (a) hbl-1::AID grown without 5-Ph-IAA. (b) Tail of a male moved to auxin at 18hr, this individual also had 12 total seam cells. (c) Tail of a male moved to auxin at 20hr, this individual also had 13 total seam cells. (d) Tail of a male moved to auxin at 22hr, this individual also had 16 total seam cells. Seam cell nuclei and cell junctions visualized with SCM::GFP and ajm-1::GFP, respectively. Yellow arrowheads indicate ray precursor lineages. Animals are oriented anterior to the left, dorsal side up.

### *lin-41* acts during the L3 stage

*lin-41* has multiple roles in the animal including the control of developmental timing (Slack et al., 2000; Spike et al., 2014). *lin-41* null alleles are sterile, whereas loss-of-function alleles cause a precocious phenotype where seam cells differentiate precociously, producing adult alae in the L4, but seam cell counts are normal. Again, it has not been specifically addressed whether *lin-41* loss-of-function mutants skip L3 or L4 stage events.

When *lin-41::AID* animals were grown continuously on 5-Ph-IAA, short precocious alae patches (less than 50% expressivity) were observed in 38% of the animals (Fig. 12A). Animals transferred to 5-Ph-IAA at 2-hour intervals from 16 to 36 hours displayed precocious alae patches (Fig. 12A). No precocious alae was observed when animals were transferred at 38 hours or later, indicating that *lin-41*’s auxin-sensitive period ends between 36 and 38 hours of postembryonic development (Fig. 12A). L3 seam cell divisions were observed at 36 hours in development and the L3 molt around 40 hours, so, auxin sensitivity for *lin-41* ended during the L3 stage.

**Fig. 12.**
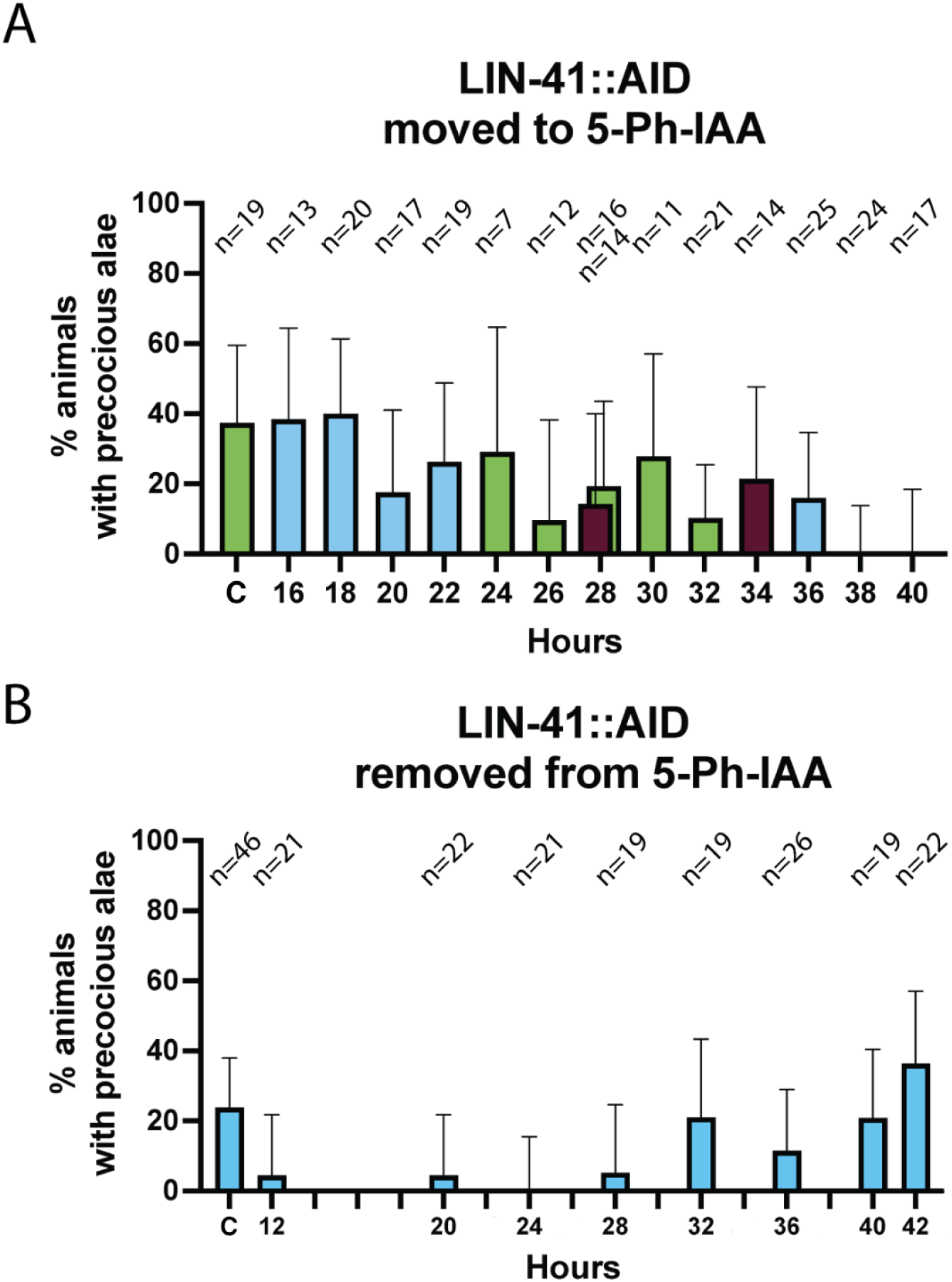
lin-41 acts primarily during the L3 stage. (A) Percent of animals with precocious alae in lin-41::AID animals transferred to 5-Ph-IAA at the indicated times after synchronization. Different bar colors indicate different groups of synchronized animals. (B) Percent of animals with precocious alae in lin-41::AID animals removed from 5-Ph-IAA at the indicated times after synchronization. ‘C’ indicates animals grown continuously on 5-Ph-IAA. Error bars indicate 95% CIs. Different experimental groups are not indicated in panel B. The total number of observed animals at each time point is indicated above the bars.

Interestingly, the fraction of animals with precocious alae patches was significantly higher in animals transferred at 16 and 18 hours than later (*p* <0.05 comparing the 16/18 hours group with the 20-40 hours group, by Fisher’s exact test). We have no explanation for this effect, but it suggests a somewhat increased sensitivity of the heterochronic pathway to *lin-41* reduction during early postembryonic stages, possibly similar to *lin-28*’s feedback loop mentioned above.

*lin-41::AID* animals moved away from 5-Ph-IAA from 32 hours and later showed percentages of precocious alae patches comparable to animals grown continuously on the auxin analog (Fig. 12B). Surprisingly, removing *lin-41::AID* animals from the auxin-analog at 12, 20, and 28 hours resulted in a small amount of precocious alae, again indicating a lasting effect of *lin-41* reduction during the L1 and L2 stages. Otherwise, these data suggest that *lin-41* acts primarily during the L3 stage.

### *lin-41* activity deficiency *likely causes skipping of L4* developmental events

We observed that male *lin-41::AID* animals grown on auxin had a precocious tail tip retraction and disrupted morphogenesis of rays and fan, as has been previously described for loss-of-function mutants (Del Rio-Albrechtsen et al., 2006). However, all ray precursor cells were produced, indicating that the proliferative cell divisions that produce them occurred normally (Fig. 13). So, in contrast with *hbl-1*, reduction of *lin-41* using the AID system did not cause skipping of L3 events. We therefore interpret the precocious differentiation of the hypodermal cells in both male and hermaphrodite as reflecting the skipping of L4-specific cell fates when *lin-41* activity is reduced.

**Fig. 13.**
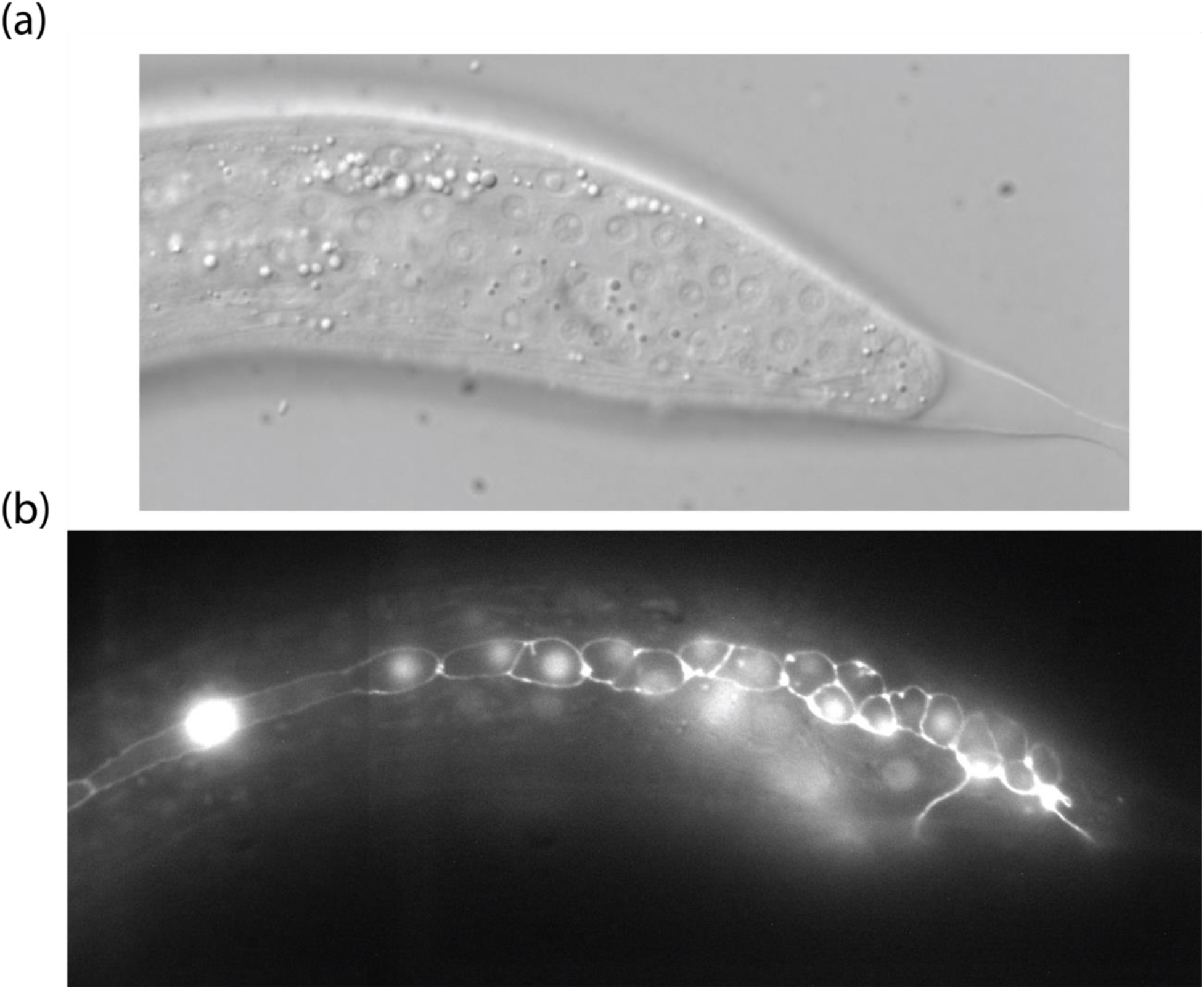
lin-41 activity reduction causes premature tail tip retraction but no change in the number of ray precursor cells. (a) DIC micrograph of lin-41::AID male grown on 5-Ph-IAA observed at 40 hr; (b) Fluorescent microscopy of the same tail showing seam cell nuclei and cell junctions visualized with SCM::GFP and ajm-1::GFP, respectively. The number of ray precursor cells appears normal. The animal is oriented anterior to the left, dorsal side up.

## Discussion

Previously, time-of-action data existed for only one heterochronic gene, *lin-14*, which was obtained using temperature-sensitive alleles (Ambros and Horvitz, 1987). The times of action of other core heterochronic genes were only suggested by indirect evidence: the earliest developmental events they affected and their expression patterns. *lin-28*, for example is expressed from the start of postembryonic development through the L2 and L3, but the first developmental it controls is at the beginning of the L2, about the time it begins to be downregulated (Ambros and Horvitz, 1984; Moss et al., 1997). In order to better understand the mechanistic relationships among the heterochronic genes, it is important to know more precisely when their activities are needed to specify stage-specific cell fates.

We found that two other heterochronic genes, *lin-28* and *hbl-1* each have two activities that, like *lin-14*’s, are separated in time (Fig. 14). Relative to events they control, both *lin-28* and *hbl-1* appear to be required just prior to or concurrent with the L2-specific symmetric seam cell division which they control, and relative to each other, *lin-28* and *hbl-1* act essentially simultaneously. By contrast, *lin-14*’s second activity (*lin-14b*) ends well before those of *lin-28* and *hbl-1* despite the fact that they control the same developmental events. Developmental timing regulators may act directly on the events they control, or they may act in advance of those events, perhaps by setting up a regulatory condition or acting through a series of molecular events that causes a delay. We know that *lin-14* controls L2 cell fates by transcriptionally repressing the expression of microRNA genes that repress *lin-28* and *hbl-1* (Tsialikas et al., 2017). Possibly, this regulation is responsible for the separation in the genes’ times of action: given that down-regulation of *lin-14* would cause transcriptional activation of miRNA genes, it may take time for that transcriptional activation to lead to miRNA-mediated repression of *lin-28* and *hbl-1* in time to have an effect on L2 fates.

**Fig. 14.**
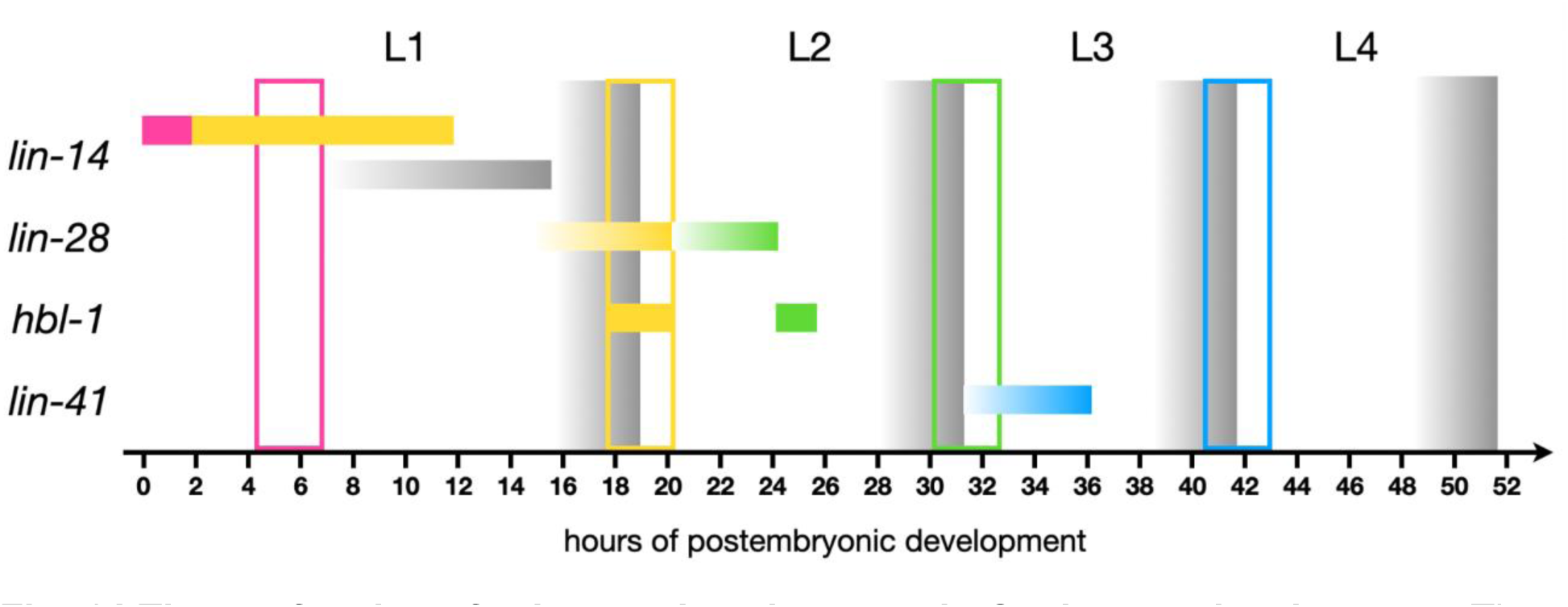
Times of actions for heterochronic genes in C. elegans development. The lethargus periods preceding and including the molts are indicated by gray gradients. Colored vertical boxes indicate approximate periods of stage-specific seam cell divisions, and colored bars indicate the genes’ auxin-sensitive periods affecting stage-specific cell fates: pink: L1 fates; yellow: L2 fates; green: L3 fates; blue: L4 fates. Gray bar indicates lin-14’s auxin-sensitive period for intestinal nuclear divisions. The bars indicate the periods when most animals showed a strong loss-of-function phenotype. The gradient in the bars is to indicate that we did not determine precisely the start time of the auxin-sensitive period.

Relative to the L3 events they control, *lin-28* and *hbl-1* are more like *lin-14b* in they act well in advance of those events: from the early-to mid-L2 to control events that happen in the early L3 (Fig. 14). Relative to each other, these genes appear staggered in time, with *lin-28* acting just prior to *hbl-1*. Genetic analysis suggests indeed that *lin-28* acts indirectly through *hbl-1* to control cell fates (Vadla et al, 2012; Ilbay and Ambros, 2019; Ilbay et al., 2021). Thus, whereas the first activities of *lin-14*, *lin-28*, and *hbl-1* act just prior to or even concurrent with the events, the second of each gene’s two activities that acts well in advance of the events it controls (Fig. 14; Ambros and Horvitz, 1987).

We identified a single activity of *lin-41* that acts in the early-to mid-L3 to control events of the L4 (Fig. 14). It is possible that yet unidentified series of molecular events intervenes between *lin-41*’s time of action and the L4 events it controls. Because null alleles of *lin-41* are sterile, we know that our *lin-41::AID* allele did not fully reduce *lin-41* activity because these animals were fertile when grown on the auxin analog (data not shown). Because we did not reduce *lin-41* activity, we cannot not be confident we have detected all of its role in the heterochronic pathway. Specifically it is theoretically possible that *lin-41*, like the other genes, has two activities, one of which regulates the L3, as previously was speculated (Vadla et al., 2012). Therefore, it may be due to the inability of our *lin-41::AID* allele which did not fully reduce *lin-41* activity that we may not have detected any activity regulating L3 events that *lin-41* might have.

There are other limitations of the AID system for precise timing of these genes. First, we know that the degron system behaves differently with different proteins (Hills-Muckey *et al*. 2021). Differences in auxin-sensitive periods may be true differences or due to differences in the degradation and activity kinetics of these factors (For instance, the HLB-1 protein may take longer to degrade than LIN-28). So, a truely precise comparison between the times of activities of the heterochronic genes is not possible. Another limitation, which is well known, is that the degradation and reactivation occur at best on the order of an hour or two, even when they occur robustly (Zhang *et al*. 2015; Ashley et al., 2021). This means that estimations of start and end times of the heterochronic genes’ periods of action cannot be precise within hours.

We unexpectedly uncovered a phenomenon that may reflect a feature of heterochronic gene hierarchy. Although restoration of activity usually occurs when auxin is removed, we found that reduction of *lin-28* activity was irreversible, and consequently, were unable to define the start of its auxin-sensitive period. This occurred even when removal from auxin was 24 hours or more from the auxin-sensitive period, theoretically enough time for the protein to recover (data not shown; Zhang et al., 2015). The *lin-28* gene has the interesting property of being both a negative regulator and a target of let-7-family microRNAs (Vadla et al. 2012; Tsialikas et al., 2017). It is possible that the auxin-induced down-regulation of *lin-28* leads to an upregulation of let-7 and therefore irreversible repression of *lin-28* expression.

Interestingly, we also observed a similar phenomenon for *lin-41*. We found *lin-41* activity was required mostly between 32-38 hours of development to control when adult alae appeared. But the precocious alae phenotype is more penetrant in animals grown continuously on 5-Ph-IAA or moved to the auxin analog before 20 hours, suggesting that *lin-41* might have some activity during the L1 and L2 stages. It is possible that *lin-41*, like *lin-28*, inhibits a negative regulator of itself, which, when derepressed prior to *lin-41*’s auxin-sensitive period, can lead to further reduction in *lin-41* activity.

Our study has placed the heterochronic gene activities into a timeline of postembryonic development relative to one another and to the developmental events whose timing they control. Further mechanistic understanding will be needed to understand how these genes control these specific events and how they influence the times of action of each other in the heterochronic gene hierarchy.

## Materials and Methods

### Strains and culture conditions

Nematodes were grown at 20°C on standard NGM plates seeded with *E. coli* AMA1004 unless otherwise indicated. Males for analysis of male tail seam cells were produced by growing strains on bacteria containing the RNAi plasmid pLT651 (Timmons et al., 2014).

### Strains used

RG733 (*wIs78* [pDP#MM016B (*unc-119*) + pJS191 (*ajm-1::GFP* + pMF1(*SCM::GFP*) + F58E10]) (wild type for this study),

HML1029 *cshIs140[rps-28pro::TIR1(F79G)_P2A mCherry-His-11; Cbr-unc-119(+)] LGII,* ME502 *cshIs140[rps-28pro::TIR1(F79G)_P2A mCherry-His-11; Cbr-unc-119(+)] LGII; hbl-1(aeIs8[hbl-1::AID]); wIs78,*

ME504 *cshIs140[rps-28pro::TIR1(F79G)_P2A mCherry-His-11; Cbr-unc-119(+)] LGII; lin-41(aeIs10[lin-41::AID]); wIs78*,

ME507 *cshIs140[rps-28pro::TIR1(F79G)_P2A mCherry-His-11; Cbr-unc-119(+)] LGII; lin-14(aeIs5[lin-14::AID]); wIs78*.

ME506 *cshIs140[rps-28pro::TIR-1(F79G)_P2A mCherry-His-11; Cbunc-119(+)] chrom II; lin-28(aeIs6[lin-28::AID]); wIs78*

### Synchronization at hatching

To generate developmentally synchronized populations of animals, adults filled with eggs were washed from crowded plates into 15ml tubes, spinned down, the excess liquid then was removed leaving a worm pellet. 500-1000µl of household bleach solution was added to the tube and vortexed every 2 minutes until cuticles of animals were dissolved enough to release eggs. Tubes then were filled with sterilized distilled water to slow the bleach activity and centrifuged at 400g for 3 minutes, then the supernatant was discarded and eggs were washed twice with sterilized distilled water and then twice with M9. Then eggs were transferred into M9 and left on a shaker for 20-48 hours at room temperature. Newly hatched larvae will not initiate L1 development without a food source. Then the liquid with larvae was centrifuged to concentrate larvae and they were transferred to plates with food, either with or without 5-Ph-IAA. The 0 time point (0 hours of postembryonic development) was when the animals were placed on food.

### Microscopy

Animals were examined using DIC and fluorescence microscopy on a Zeiss Axioplan2 microscope with Zeiss objectives: Plan-NEOFLUAR 5x, Plan-NEOFLUAR 16x, Plan-NEOFLUAR 63x, alpha Plan-FLUAR 100x. Images were acquired using AxioCam with AxioVision software.

### CRISPR/Cas9

We followed general protocols described in Paix *et al*. 2017. gRNA was synthesized from PCR-amplified templates using Invitrogen™ MEGAshortscript™ T7 Transcription Kit (Catalog# AM1354) and purified with Invitrogen™ MEGAclear™ Transcription Clean-Up Kit (Catalog# AM1908). Purified gRNAs were mixed with Cas9 (EnGen^®^ Spy Cas9 NLS, Catalog# M0646T) and used in microinjections. Typical concentrations of the components in the injection mix: target gRNA (up to 100ng/µl), Cas9 (250 ng/µl), co-injection *dpy-10* gRNA (100 ng/µl).

We inserted minimal degron sequence (Zhang *et al*. 2015; Morawska and Ulrich, 2013) at the 3’ end of the open reading frame in *lin-14*, *lin-28*, and *hbl-1* and at the 5’ end of the open reading frame in *lin-41*.

To make insertions, a hybrid dsDNA repair template was used as described in (Dokshin *et al*. 2018). Repair templates were melted and cooled before injections (Ghanta and Mello, 2020). The repair template then was added to the CRISPR mix to the final concentration of 100-500 ng/µl of DNA.

Progeny that had roller phenotypes was isolated and then, plates were screened for AID insertion. AID alleles were outcrossed 3x times.

### Auxin-inducible degron system

We used a modified auxin-inducible system with *TIR1(F79G)*, using 5-Ph-IAA as the auxin analog (Zhang *et al*. 2015; Hills-Muckey *et al*. 2021). To make 5-Ph-IAA-containing plates, 50µmol of 5-Ph-IAA that was spread on standard NGM plates to approximately 0.005 µM concentration in the agar prior to seeding.

Synchronized animals were placed either onto NGM plates without 5-Ph-IAA, then transferred to 5-Ph-IAA at the indicated time point, or onto plates with 5-Ph-IAA, then transferred to NGM plates without 5-Ph-IAA at the indicated time point.

Because it was not practical to cover all time points in a single experiment, experiments were performed for a subset of all the time points required to define auxin-sensitive periods. Table S1 lists each experiment and the strains and time points covered in each, which are also indicated in the figures.

### Plots and statistics

Data was analyzed using Prism software. *P*-values were calculated using unpaired Welch’s t-tests for absolute values (seam cell and intestinal nuclei count averages) and Fisher’s exact tests for fractions (percents). Error bars in plots indicate 95% CI, asterisks indicate the following: * *p* ≤ 0.05, ** *p* ≤ 0.01, *** *p* ≤ 0.001, **** *p* ≤ 0.0001, ns - not significant (*p* > 0.05).

## Acknowledgements

We would like to thank members of Ellis lab for valuable comments on the manuscript, Ronald Ellis for comments, advice and strains, Chris Hammell for TIR1 strains, and plasmid. We thank WormBase for genetic sequences and WormAtlas for reference information on nematode biology and development. Some strains were provided by the CGC, which is funded by NIH Office of Research Infrastructure Programs (P40 OD010440).

## Data availability Statement

Strains are available upon request.

## Supplementary information

**Table S1.** List of experiments. Strains in each experiment were synchronized at the same time and transfers were carried out at similar time points.

| Expt. number | Strains synchronized | Direction of transfers | Hours after synchronization |
| --- | --- | --- | --- |
| 1 | *lin-14::AID, lin-28::AID* | To 5-Ph-IAA | 2, 4, 6, 8, 10 |
| 2 | *lin-14::AID, lin-28::AID* | To 5-Ph-IAA | 12, 14, 16, 18, 20 |
| 3 | *lin-14::AID* | To 5-Ph-IAA | 8, 10 |
| 4 | *lin-28::AID, hbl-1::AID* | To 5-Ph-IAA | 14, 16, 18, 20, 22, 24 |
| 5 | *lin-28::AID, hbl-1::AID* | To 5-Ph-IAA | 24, 26, 28, 30, 32 |
| 6 | *lin-14::AID* | To 5-Ph-IAA | 13, 14, 16, 18, 20, 22 |
| 7 | *hbl-1::AID, lin-41::AID* | To 5-Ph-IAA | 24, 26, 28, 30, 32 |
| 8 | *hbl-1::AID, lin-41::AID* | To 5-Ph-IAA, Off 5-Ph-IAA | 12, 16, 18, 20, 22, 36, 38, 40 |
| 9 | *hbl-1::AID, lin-41::AID* | To 5-Ph-IAA, Off 5-Ph-IAA | 8, 24, 28, 32, 34 |
| 10 | *hbl-1::AID* | To 5-Ph-IAA, Off 5-Ph-IAA | 14, 16, 18, 20, 22, 24 |
| 11 | *hbl-1::AID* | To 5-Ph-IAA, Off 5-Ph-IAA | 24, 26, 28, 30 |
| 12 | *lin-14::AID* | To 5-Ph-IAA, Off 5-Ph-IAA | 1, 2, 4, 6, 8 |
| 13 | *lin-41::AID* | To 5-Ph-IAA | 36, 38, 40, 42, 44 |
| 14 | *lin-41::AID* | Off 5-Ph-IAA | 40, 42 |

**Fig. S1.**
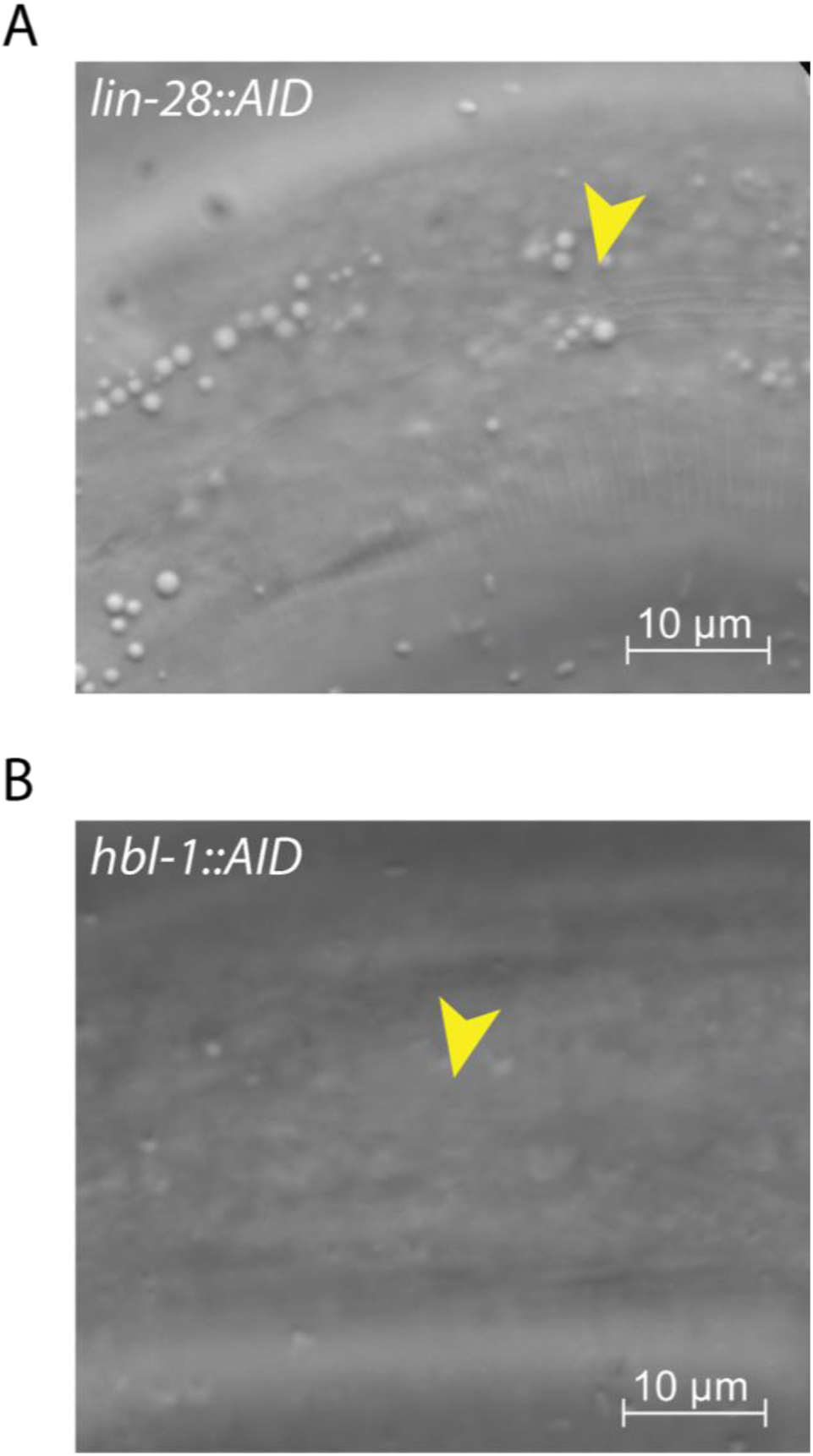
Precocious alae appearances differ in lin-28::AID and hbl-1::AID animals on 5-Ph-IAA. DIC microphotographs of L4 animals grown at 20°C. Animals are oriented anterior to the left, dorsal side up. (A) lin-28::AID animals develop clear alae patches when moved to 5-Ph-IAA at times close to the end of the second activity. The yellow arrowhead indicates the edge of an alae patch. (B) hbl-1::AID animals develop thin and transparent alae when moved to 5-Ph-IAA at times close to the end of the second activity, the edges of alae patches are hard to discern. The yellow arrowhead indicates alae.

